# Functional integrity of the SEL1L-HRD1 complex is critical for ERAD and organismal viability

**DOI:** 10.1101/2025.08.01.668162

**Authors:** Xiawei Zhang, Liangguang Leo Lin, Linxiu Pan, Xiaoqiong Wei, Huilun Helen Wang, Zexin Jason Li, Ling Qi

**Affiliations:** Department of Molecular Physiology and Biological Physics, University of Virginia School of Medicine, Charlottesville, VA 22903, USA

**Keywords:** ERAD, SEL1L-HRD1 complex, Protein quality control, SEL1L disease variants, Neonatal lethality

## Abstract

The SEL1L-HRD1 complex represents the most evolutionarily conserved branch of endoplasmic reticulum-associated degradation (ERAD), with SEL1L acting as a key cofactor for the E3 ubiquitin ligase HRD1. While the physiological relevance of this complex has been increasingly recognized, whether SEL1L is strictly required for HRD1 function in mammals has remained unclear. Here, using complementary *in vivo* and *in vitro* approaches, we define the architecture and physiological significance of the mammalian SEL1L-HRD1 ERAD complex. Our data demonstrate that direct binding between SEL1L and HRD1 is essential for ERAD function and neonatal survival in mice. In three knock-in mouse models harboring targeted mutations at the SEL1L-HRD1 interface, we show that the L709P variant – unlike the benign P699T mutation – results in complete neonatal lethality within 30 hours of birth, a phenotype more severe than that of the partially lethal S658P variant. Mechanistically, the L709P mutation abolishes SEL1L-HRD1 interaction, disrupting substrate engagement and impairing recruitment of the E2 enzyme UBE2J1, leading to the accumulation and aggregation of misfolded proteins in the ER. Notably, these defects can be partially rescued by HRD1 overexpression, echoing findings from yeast. Together, our results provide definitive evidence that the SEL1L-HRD1 interaction is essential for ERAD activity and neonatal viability in mammals, resolving a long-standing question in ERAD biology and identifying a new therapeutic strategy for modulating ERAD activity in humans.

**SIGNIFICANCE STATEMENT:** The endoplasmic reticulum-associated degradation (ERAD) pathway is essential for maintaining cellular proteostasis and organismal health, yet its molecular regulation in mammals remains incompletely defined. The SEL1L-HRD1 complex constitutes the central axis of this conserved pathway, but whether SEL1L is required for HRD1 function *in vivo* has remained unresolved. Here, using genetic mouse models and biochemical analyses, we demonstrate that the physical interaction between SEL1L and HRD1 is essential for ERAD activity and neonatal viability. These findings resolve a long-standing question in ERAD biology, provide new insights into the structural and functional organization of the mammalian ERAD machinery, and highlight the SEL1L-HRD1 interface as a potential therapeutic target for modulating ERAD activity in diseases.

## MAIN TEXT

In eukaryotic cells, approximately one-third of nascent proteins enter the endoplasmic reticulum (ER), where they fold and mature into functional conformations (1, 2). Misfolded proteins – arising from genetic mutations or folding inefficiencies (3) – are eliminated by ER-associated degradation (ERAD), a conserved quality control process that targets them for cytosolic proteasomal degradation (4-10). The most conserved ERAD branch involves the Hrd3p-Hrd1p complex in yeast and its mammalian homolog, the SEL1L-HRD1 complex (11, 12). Genetic ablation of *Sel1L* or *Hrd1* in mice, either germline or acutely in adults, leads to embryonic or premature lethality, respectively, underscoring their essential roles in development and survival (13-16). Cell-type-specific conditional deletions have further implicated SEL1L and HRD1 in diverse physiological processes, including nutrient and energy metabolism, and immune regulation (17-19). More recently, pathogenic variants in *SEL1L* and *HRD1* have been linked to human neurodevelopment disorders known as ERAD-associated neurodevelopment disorders with onset in infancy (ENDI) syndrome (20-22). Despite these advances, it remains unclear whether SEL1L and HRD1 function strictly as a complex or retain independent roles – an unresolved question with direct implications for therapeutic targeting.

Both yeast Hrd3 and mammalian SEL1L serve as scaffold for the HRD1 protein complex and contribute to HRD1 protein stability (16, 23-26). Notably, Hrd1p overexpression in yeast can bypass the requirement for Hrd3p (27-29), suggesting the possibility of Hrd3p- independent Hrd1p function. In line with this, proteomic studies in mammalian systems have revealed SEL1L-independent degradation pathways and alternative HRD1 complexes (30, 31). Conversely, SEL1L may also exert HRD1-independent roles, such as in collagen turnover (32). Thus, the physiological significance of the SEL1L-HRD1 interaction – and the extent to which these components may function independently – remains to be fully elucidated.

We previously reported that the SEL1L S658P variant, originally identified in Finnish Hounds with cerebellar ataxia (33), causes a similar neurodegenerative phenotype in mice (26). The S658 residue lies adjacent to – but not directly within – the SEL1L-HRD1 interface, and this mutation attenuates their interaction while destabilizing SEL1L protein. Consequently, the precise physiological relevance of the SEL1L-HRD1 interaction remains incompletely defined. Notably, knock-in (KI) mice homozygous for the S658P variant survive to weaning at sub-Mendelian ratios (∼ 11%), and those that do survive develop early-onset ataxia in adulthood (26).

In this study, we generated two additional KI mice harboring human SEL1L missense variants, L709P (NM_005065, c.2126T>C; p.Leu709Pro) and P699T (c.2095C>A; p.Pro699Thr) – both located within the C-terminal luminal domain of SEL1L at the SEL1L- HRD1 interface. Remarkably, homozygous L709P KI mice, but not P699T, display complete neonatal lethality within 30 hours of birth. This lethality is accompanied by a near-complete loss of SEL1L-HRD1 interaction and pronounced ERAD defects, all of which are more severe than those observed in SEL1L S658P KI mice. These findings not only demonstrate a strong correlation between SEL1L-HRD1 interaction (i.e. complex integrity), ERAD function, and organismal postnatal survival, but also uncover a structurally defined node that may serve as a therapeutic target for diseases driven by SEL1L-HRD1 ERAD dysfunction and broader ER proteostasis defects.

## RESULTS

### Generation of two KI mice carrying human SEL1L variants

Through collaboration with Baylor’s clinical genomic sequencing program, we identified two heterozygous SEL1L missense variants, P699T and L709P, in pediatric patients (Table S1). Both variants map to the highly conserved Sel1-like repeat C-terminal (SLR- C) domain of SEL1L (Fig. 1A) and involve residues that are evolutionary conserved across vertebrate species (Fig. 1B). Position-specific scoring matrix (PSSM) analysis indicated that both substitutions occur at residues under strong evolutionary constraint, suggesting that both are likely to be functional detrimental (Fig. S1A-B). Consistently, PolyPhen-2, a computational tool that predicts the potential impact of amino acid substitutions on protein structure and function (34), classified both variants as ‘possibly damaging’ (Table S1).

**Figure 1.**
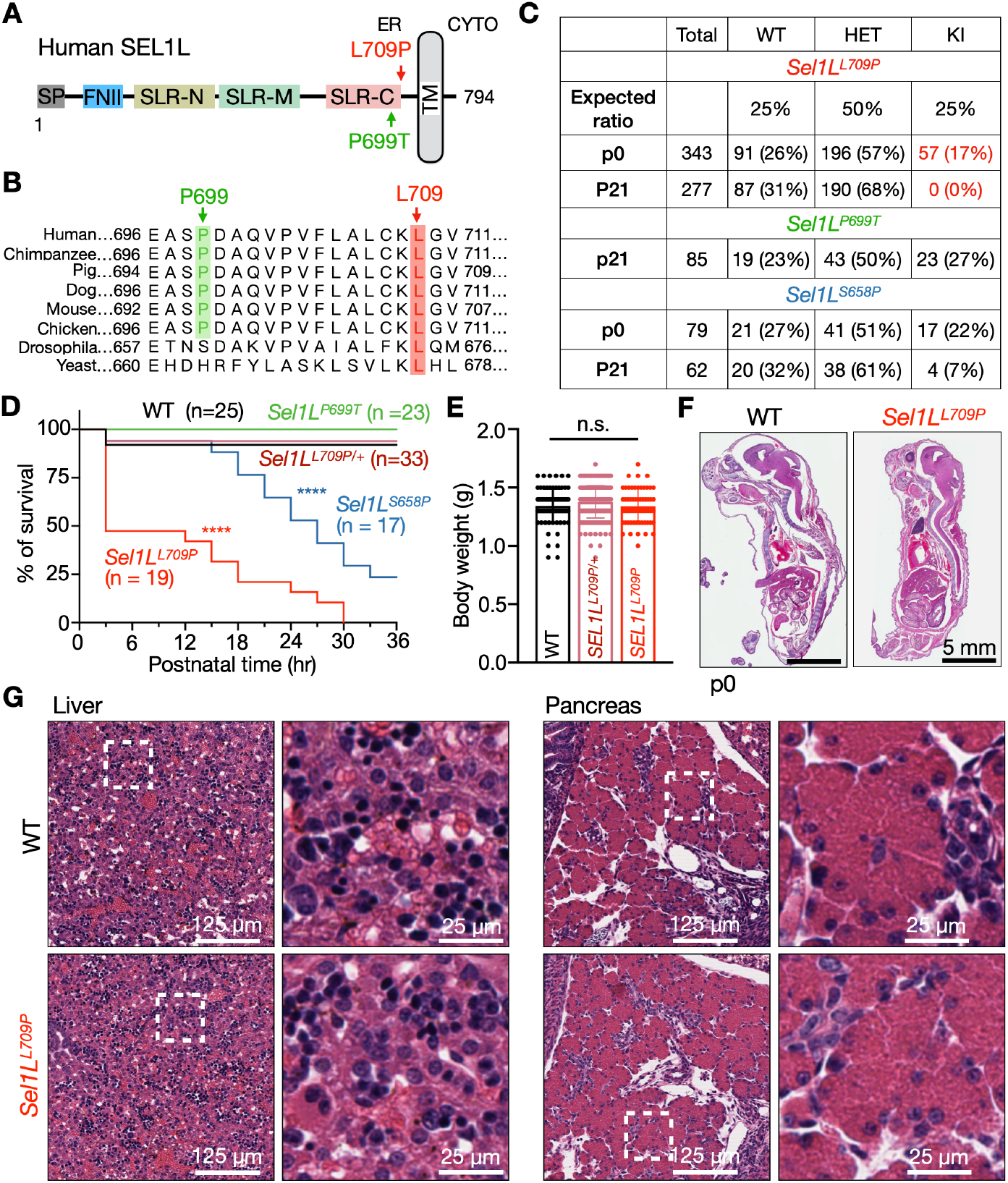
Homozygous *Sel1L*^*L709P*^ KI mice exhibit neonatal lethality. (A) Schematic diagram of the human SEL1L protein domain structure, highlighting the position of the variants. SP, signal peptide; FNII, fibronectin type II domain; SLR-N/M/C, N-, middle-, and C-terminal Sel1-like repeats; TM, transmembrane domain. (B) ClustalW sequence alignment demonstrating evolutionary conservation of residues L709 (red) and P699 (green). (C) Number and percent of different genotypes at different postnatal (p) ages. (D) Kaplan–Meier survival curves for neonates over the first 30 hours after birth. n, mouse numbers. ^****^, p < 0.0001 (comparing KI to WT mice) by Log-rank (Mantel–Cox) test. (E) Body weights of WT and KI neonates at postnatal day 0 (P0). Data, mean ± SEM; n.s., not significant by one-way ANOVA with Dunnett’s multiple comparisons test. (F-G) Hematoxylin and eosin (H&E) staining of WT and *Sel1L*^*L709P*^ KI p0 pups. Various internal organs appeared histologically normal (G). n = 3 mice per group.

To investigate functional consequences of these variants in vivo, we generated KI mice carrying the orthologous P699T or L709P mutations via CRISPR/Cas9-mediated genome editing (Fig. S1C). Two independent founder lines were established for each mutation (Fig. S1D) and maintained separately. As both lines displayed comparable phenotypes, data from each were pooled for subsequent analysis.

### The L709P variant, but not P699T, causes neonatal lethality in mice

Strikingly, while *Sel1L*^*P699T*^ mice were recovered at ∼25% of expected Mendelian ratios at weaning, no homozygous *Sel1L*^*L709P*^ mice were recovered among more than 300 genotyped pups (Fig. 1C). However, homozygous *Sel1L*^*L709P*^ pups were born at near-Mendelian ratios (∼17%) at postnatal day 0 (P0), indicating that lethality occurred after birth. These newborns appeared grossly normal (Fig. S1E) but died uniformly within 30 hours of birth, with over 50% found dead within the first 3 hours (Fig. 1D). Heterozygous *Sel1L*^*L709P/+*^ mice were born and survived to weaning at expected Mendelian ratios (Fig. 1C-D). At birth, body weights were comparable among *Sel1L*^*L709P*^, heterozygous *Sel1L*^*L709P/+*^, and wild-type (WT) littermates (Fig. 1E).

For comparison, *Sel1L*^*S658P*^ homozygotes exhibited a less severe phenotype, with ∼50% dying after 30 hours and a quarter survived beyond weaning. Histological analysis of major organs—including pancreas, liver, brain, and heart—revealed no overt developmental abnormalities in *Sel1L*^*L709P*^ neonates (Fig. 1F-G and Fig. S2). In contrast, both *Sel1L*^*L709P/+*^ heterozygotes and *Sel1L*^*P699T*^ homozygotes were phenotypically indistinguishable from WT controls (Fig. 1C-D; Fig. S1F-G). Together, these data demonstrate that the L709P variant, but not P699T, results in neonatal lethality in mice, representing a more severe phenotype than S658P.

### SEL1L L709P impairs ERAD function *in vivo* and *in vitro*

To evaluate how the SEL1L L709P mutation affects ERAD function, we analyzed protein levels of core ERAD components (SEL1L, HRD1, and OS9) using KI mouse tissues and mouse embryonic fibroblasts (MEFs). We compared livers and brain tissues from three KI lines (*Sel1L*^*L709P*^, *Sel1L*^*P699T*^, and *Sel1L*^*S658P*^) with hepatocyte-specific *Sel1L* knockout mice (*Sel1L*^*AlbCre*^) as controls (35-37). In p0 livers, SEL1L protein levels were unexpectedly elevated about twofold in *Sel1L*^*L709P*^ mice relative to WT controls (Fig. 2A and quantified in Fig. 2B), whereas they were reduced by over 50% in *Sel1L*^*S658P*^ mice. In *Sel1L*^*AlbCre*^ livers, SEL1L protein levels were similarly reduced by ∼50%, likely due to contribution from non-hepatocyte populations (35-37). Despite the increase SEL1L protein, HRD1 levels were reduced by ∼45% in *Sel1L*^*L709P*^ liver, similar to the reduction observed in *Sel1L*^*S658P*^ livers, whereas HRD1 levels remained unchanged in *Sel1L*^*AlbCre*^ livers (Fig. 2A and quantified in Fig. 2B). OS9, an ER lectin and ERAD co-factor, was elevated approximately threefold in *Sel1L*^*L709P*^ and tenfold in *Sel1L*^*AlbCre*^ livers but remained unchanged in *Sel1L*^*S658P*^ mice (Fig. 2A-B). These changes occurred independently of mRNA expression (Fig. 2C), indicating post-transcriptional regulation. Similar patterns were observed in p0 brain tissue (Fig. S3A and quantified in S3B).

**Figure 2.**
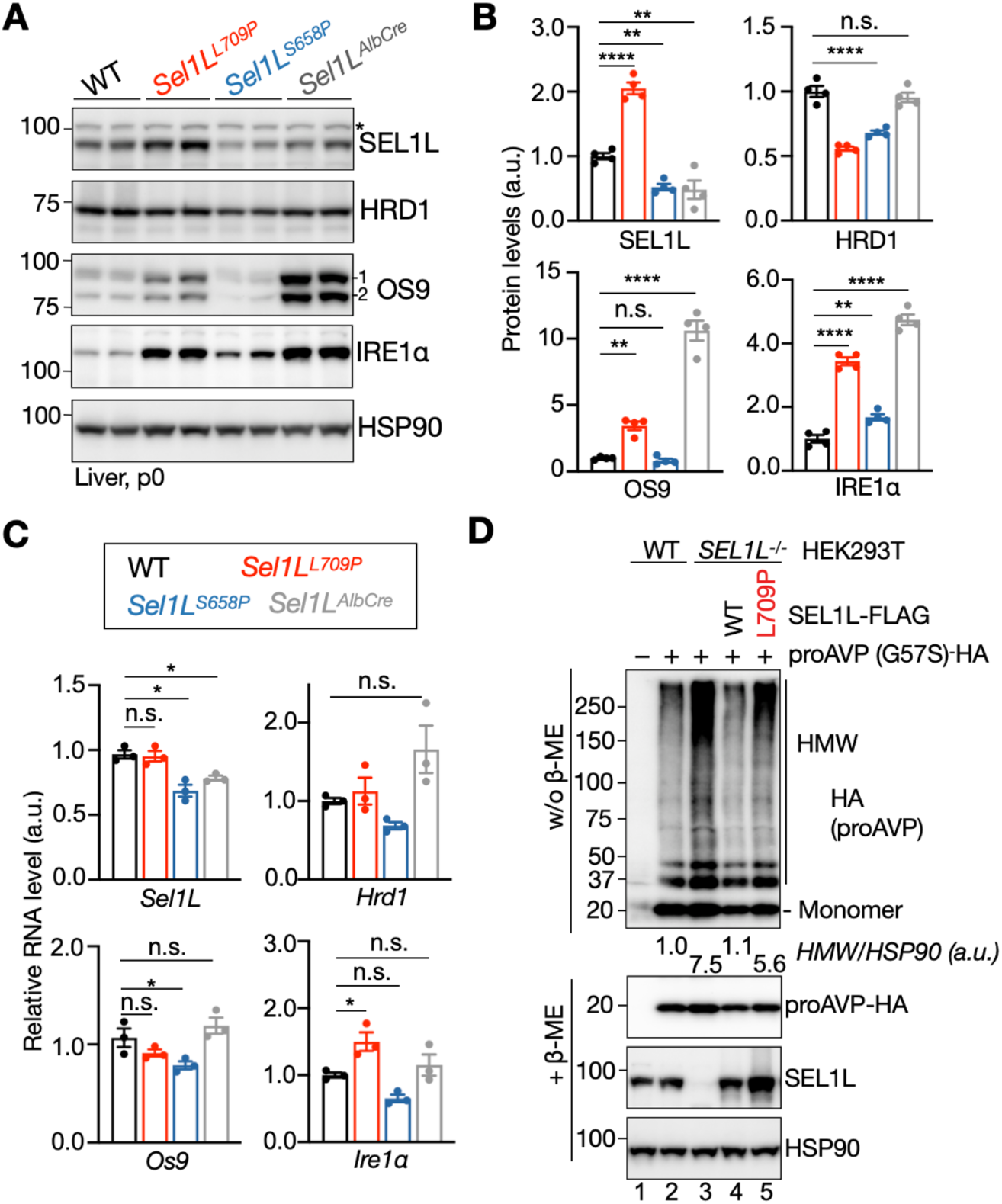
SEL1L L709P mutation impairs ERAD function *in vivo* and *in vitro*. (A-B) Immunoblot analysis of core ERAD components (SEL1L, HRD1, OS9) and the endogenous substrate IRE1α in livers from WT, *Sel1L*^*L709P*^ KI, *Sel1L*^*S658P*^ KI, and *Sel1L*^*AlbCre*^ mice (A), with quantification of protein levels shown in (B), normalized to the loading control HSP90. Non-specific bands are indicated by asterisks. Data represents four mice per group, with 2 repeats. (C) qPCR analysis of ERAD-related gene expression in livers, normalized to *L32*. Data represents three mice per group, with 2 repeats. (D) Reducing and non-reducing SDS-PAGE followed by immunoblotting to detect high-molecular- weight (HMW) aggregates of the misfolded ERAD substrate proAVP-G57S in *SEL1L*^*-/-*^ HEK293T cells expressing the indicated *Sel1L-FLAG* constructs (WT, S658P, or L709P). Quantification of HMW AVP shown below the gel represents the average of four independent experiments, normalized to the loading control HSP90. Data represents four independent experiments, with 3 repeats. Values are mean ± SEM. n.s., not significant; ^*^p < 0.05; ^**^p < 0.01; ^***^p < 0.001; ^****^p < 0.0001 by one-way ANOVA with Dunnett’s multiple comparisons test.

We next assessed ERAD component expression in MEFs from the KI lines. Inducible *Sel1L* knockout (*Sel1L*^*ERCre*^) MEFs (16) were included as controls. As in vivo, *Sel1L*^*L709P*^ MEFs showed elevated SEL1L protein (albeit not reaching statistical significance), a ∼40% reduction in HRD1, and a twofold increase in OS9 levels (Fig. S3C-D and quantification shown in Fig. S3E). These changes were comparable to those observed in *Sel1L*^*ERCre*^ MEFs (Fig. S3C-D and quantification shown in Fig. S3E). In contrast, *Sel1L*^*P699T*^ MEFs resembled WT MEFs across all markers.

To directly evaluate ERAD function, we measured protein levels of the endogenous ERAD substrate IRE1α (38) and high-molecular-weight (HMW) aggregates of the model substrate proAVP-G57S (39). IRE1α protein was elevated ∼3.5-fold in *Sel1L*^*L709P*^ livers, falling between the ∼4.7-fold increase observed in *Sel1L*^*AlbCre*^ mice and ∼1.7-fold increase seen in *Sel1L*^*S658P*^ mice (Fig. 2A and quantified in Fig. 2B). *Ire1a* mRNA was elevated by ∼ 40% in *Sel1L*^*L709P*^ livers, but unchanged in both *Sel1L*^*S658P*^ and *Sel1L*^*AlbCre*^ livers (Fig. 2C), indicating a predominantly post-transcriptional accumulation. These findings were confirmed in brain tissue (Fig. S3A and quantified in S3B) and MEFs (Fig. S3C-D and quantified in S3E).

We next assessed disulfide-linked aggregation of proAVP-G57S-HA In *SEL1L*^*-/-*^ HEK293T cells. While overexpression of WT SEL1L significantly reduced aggregation, the L709P variant led to only a modest decrease (lanes 3-5, Fig. 2D). Collectively, these data demonstrate that the SEL1L L709P mutation severely impairs ERAD function despite maintaining SEL1L protein levels.

### SEL1L L709P expression is associated with subtle unfolded protein response (UPR) activation

Previous studies showed that SEL1L-HRD1 ERAD deficiency leads to a subtle activation of UPR, likely due to cellular adaptation involving ER dilation, increased expression of various ER chaperones and activation of ER-phagy (17, 19, 40, 41). To assess whether the *Sel1L*^*L709P*^ mutation triggers a similar response, we examined two key UPR branches in P0 tissues: IRE1α and PERK. In *Sel1L*^*L709P*^ livers, while total IRE1α protein was increased ∼3.5-fold (Fig. 2A-B), the ratio of phosphorylation IRE1α to total IRE1α was not significantly increased (Fig. 3A; quantified in Fig. 3B). Consistently, *Xbp1* mRNA splicing, a downstream output of IRE1α activation was largely unchanged (Fig. 3C-D). BiP (GRP78) expression was modestly elevated (∼1.7-fold) (Fig. 3A and quantified in Fig. 3B). For the PERK pathway, we observed a mild increase in PERK phosphorylation and a ∼2-fold increase in the phosphorylation of its downstream effector eIF2α (Fig. 3F-G). The magnitude of BiP and PERK/eIF2α phosphorylation was comparable in *Sel1L*^*AlbCre*^ livers and greater than in *Sel1L*^*S658P*^ livers (Fig. 3).

**Figure 3.**
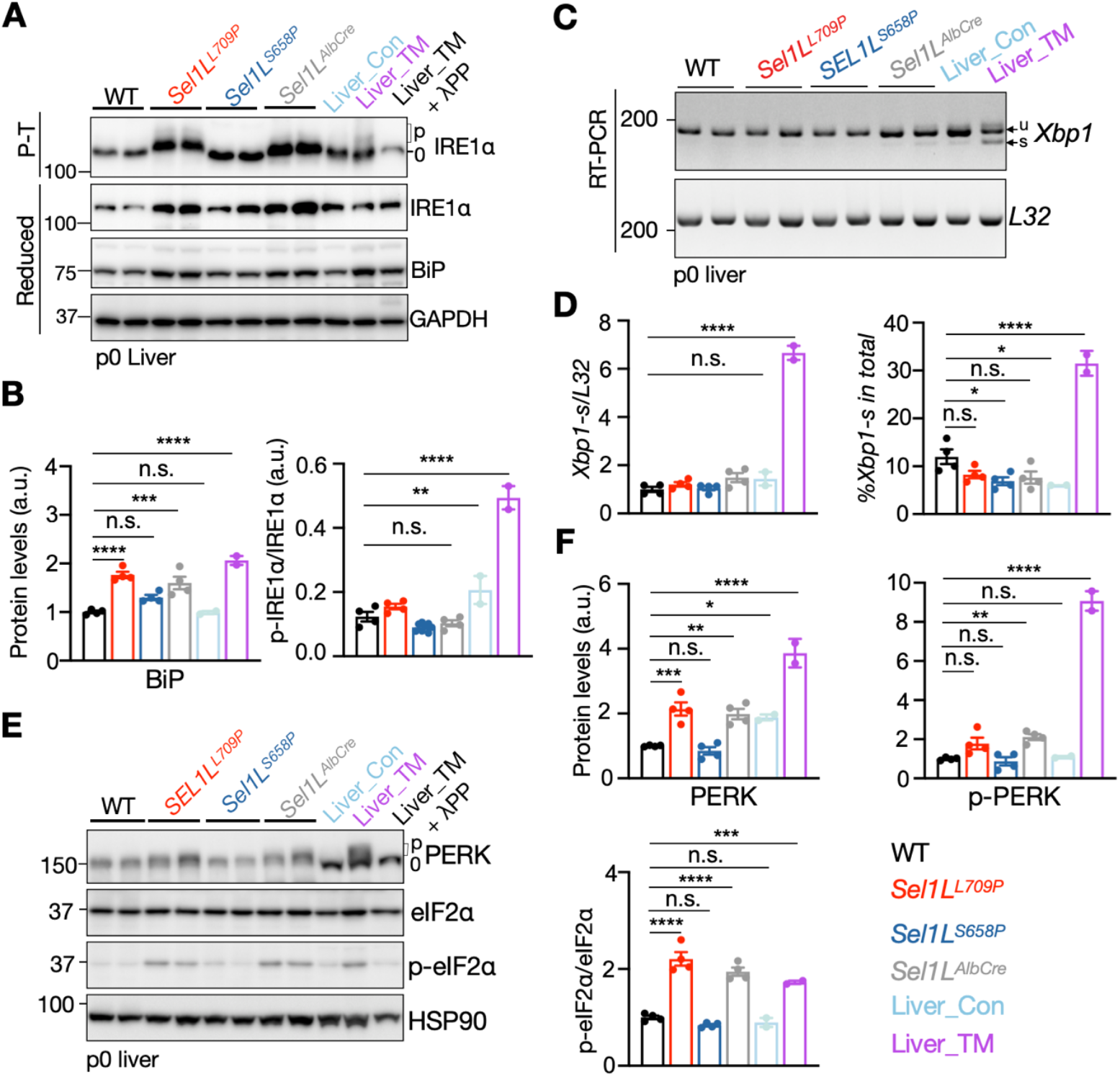
*Sel1L*^*L709P*^ expression is associated with mild ER stress *in vivo*. (A-B) Phos-tag gel analysis of phosphorylated IRE1α in liver tissues from WT, *Sel1L*^*L709P*^, and *Sel1L*^*S658P*^ KI, *Sel1L*^*AlbCre*^ mice and control liver tissues (Liver_Con: treated with PBS; Liver_TM: treated with TM) (A). λPP treatment was used to show IRE1α phosphorylation. Total IRE1α and BiP were assessed using the standard SDS-PAGE. Quantification of BiP protein level normalized to GAPDH and of p-IRE1α/IRE1α shown in (B). Data represents four mice per group, with 3 repeats. (C–D) RT-PCR analysis of *Xbp1* mRNA splicing in livers (C), with quantification shown in (E). Data represents four mice per group, with 2 repeats. (E-F) Immunoblot analysis of PERK, total and phosphorylated eIF2α (p-eIF2α) in livers (E), with quantification of protein levels shown in (F), normalized to HSP90. Data represents four mice per group, with 3 repeats. Values are shown as mean ± SEM. n.s., not significant; ^*^p < 0.05; ^**^p < 0.01; ^***^p < 0.001; ^****^p < 0.0001 using one-way ANOVA with Dunnett’s multiple comparisons test.

### SEL1L L709P induces mild ER dilation in vivo

Transmission electron microscopy (TEM) of liver and pancreatic tissues revealed structural ER alterations. While WT and *Sel1L*^*L709P*^ hepatocytes retained sheet-like ER morphology with abundant ribosomes (arrows), mild ER dilation was apparent in *Sel1L*^*L709P*^ cells (asterisk, Fig. 4A). In pancreatic acinar cells, zymogen granules appeared normal, but the ER was visibly swollen in *Sel1L*^*L709P*^ mice (asterisk, Fig. 4B). These ultrastructural changes support that SEL1L L709P is associated with a subtle UPR *in vivo* with mild ER dilation.

**Figure 4.**
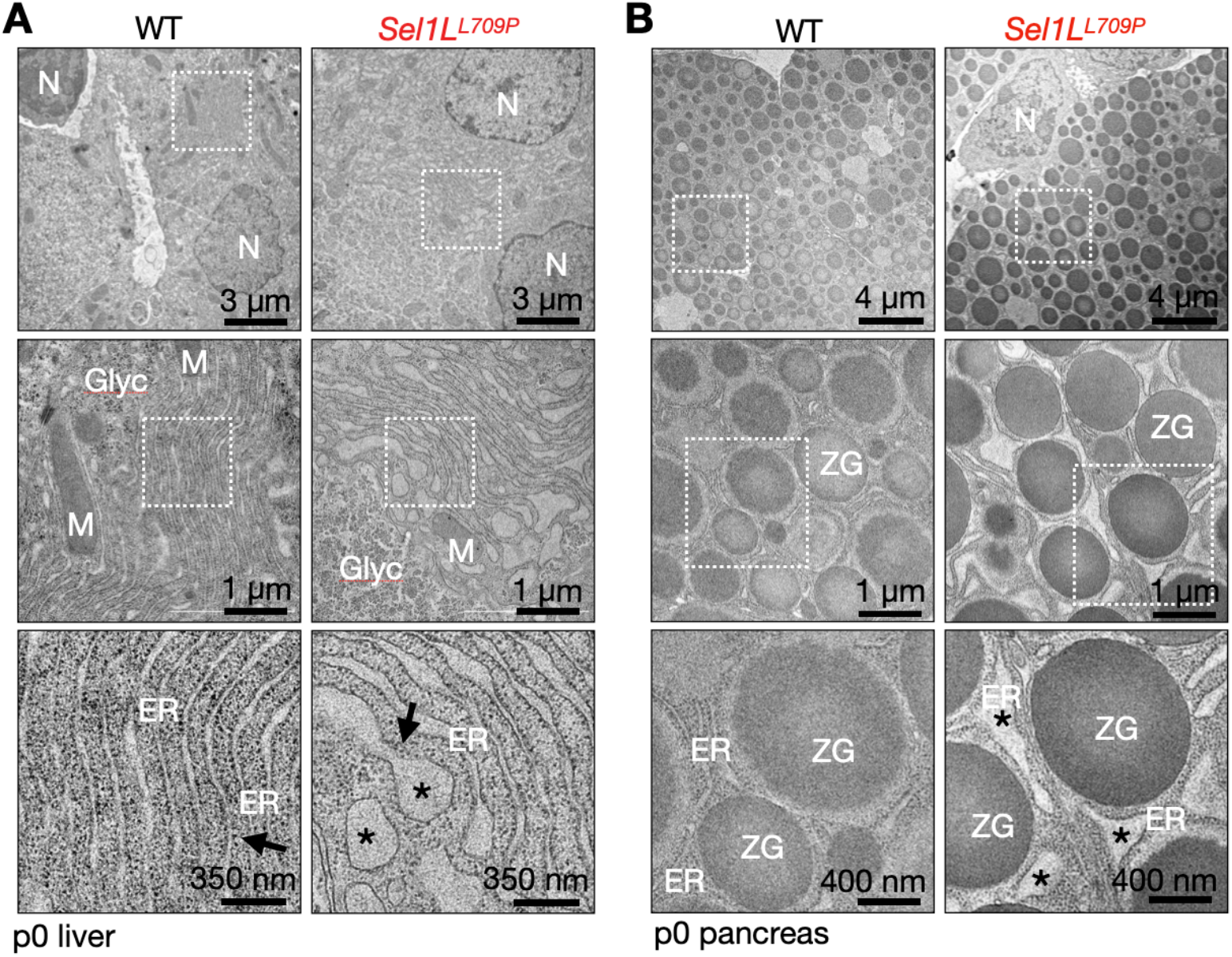
SEL1L L709P mutation causes mild ER dilation in liver and pancreas. (A) Representative transmission electron microscopy (TEM) images of livers from P0 WT and *Sel1L*^*L709P*^ mice. N, nucleus; M, mitochondria; ER, endoplasmic reticulum; Glyc, glycogen. Asterisk, dilated ER; Arrows, the ribosomes. n = 3 mice with 30-40 cells per mouse. (B) Representative TEM images of pancreatic acinar cells. ZG, Zymogen granules; N, nucleus; M, mitochondria; ER, endoplasmic reticulum. Asterisk, dilated ER. n = 3 mice with 20-30 cells per mouse.

### SEL1L L709P abolishes SEL1L-HRD1 interaction

Our recent cryo-EM structure of the mammalian OS9-SEL1L-HRD1 complex revealed a dimeric arrangement (Fig. 5A) distinct from its yeast counterpart (42-44). L709 resides within the amphipathic helix (APH) of SEL1L, making direct contact the TM1-TM2 loop of HRD1, whereas P699 lies within the adjacent loop region (Fig. 5B). Structural modeling suggests that substituting L709 with proline may disrupt this interaction due to local distortion of the APH. S658 resides in a separate helix that is near, but not directly within, the SEL1L-HRD1 interfaces (Fig. 5B).

**Figure 5.**
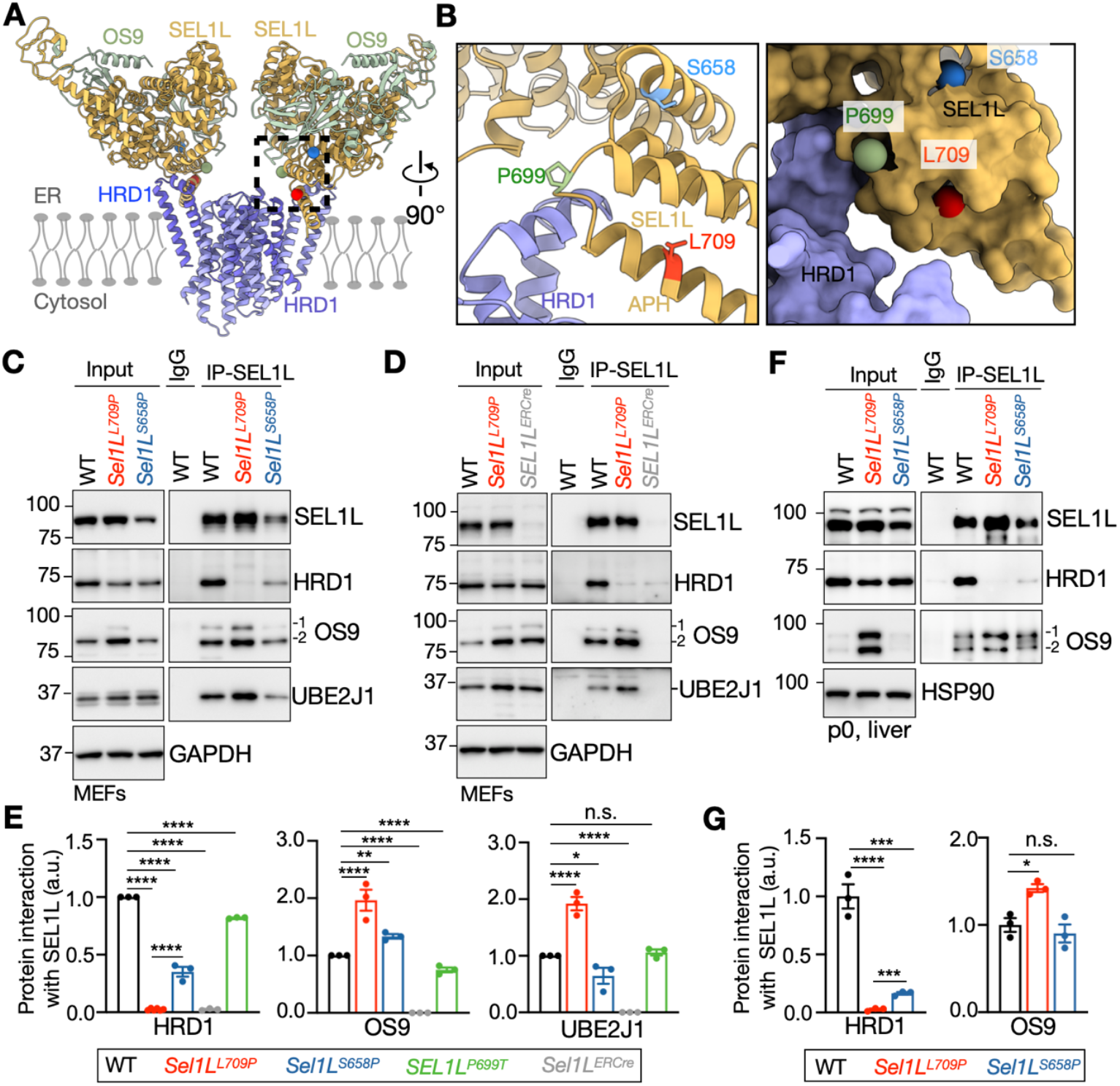
SEL1L L709P mutation abolishes SEL1L-HRD1 interaction *in vivo* and *in vitro*. (A-B) Dimeric OS9-SEL1L-HRD1 complex structure (A). A close-up view highlights the SEL1L- HRD1 interface with the location of three variants shown (B). APH, amphipathic helix of SEL1L. (C–E) Co-immunoprecipitation (Co-IP) of endogenous SEL1L from MEFs derived from WT, *Sel1L*^*S658P*^, *Sel1L*^*L709P*^ and *Sel1L*^*ERCre*^ mice, followed by immunoblotting for ERAD components (C- D), with quantification shown in (E); GAPDH serves as a loading control. Data represents the MEFs from n = 3 mice per group, with 2 repeats. (F-G) Co-IP of endogenous SEL1L from livers of p0 neonates (WT, *Sel1L*^*S658P*^, and *Sel1L*^*L709P*^ mice), followed by western blot analysis of ERAD components (F), with quantification shown in (G). HSP90 serves as a loading control. Data represents three mice per group, with 2 repeats. Values, mean ± SEM. n.s., not significant; ^*^p < 0.05; ^**^p < 0.01; ^***^p < 0.001; ^****^p < 0.0001 using one-way ANOVA with Dunnett’s multiple comparisons test.

Co-immunoprecipitation assays in *Sel1L*^*L709P*^ MEFs confirmed a near-complete loss of HRD1 binding, while interactions with the E2 enzyme UBE2J1 (38) and the lectin OS9 were preserved or mildly increased (Fig. 5C-D and S4A; quantified in Fig. 5E). In contrast, the P699T variant did not affect HRD1 binding, and S658P reduced it by ∼60% (Fig. 5C-D and S4A, quantified in Fig. 5E). Similar results were observed in p0 livers, where the L709P mutation led to a near-total (∼100%) loss of HRD1 binding – significantly more severe than the ∼ 80% reduction observed with S658P (Fig. 5F, quantified in Fig. 5G). The interaction between SEL1L and OS9 were not affected by either SEL1L variant (Fig. 5C-D, quantified in Fig. 5E). Reverse immunoprecipitation using HRD1 confirmed the loss of SEL1L-HRD1 binding in *Sel1L*^*L709P*^ MEFs (Fig. S4B and quantified in Fig. S4C).

To mechanistically explain the disruption caused by SEL1L L709P mutation, we next tested the importance of the APH in ERAD by deleting residue 699-720 (ΔAPH) (Fig. S4D). In *SEL1L*^*-/-*^ HEK293T cells, while overexpression of WT SEL1L, P699T or S658P reduced proAVP-G57S aggregation (lanes 3, 5, and 6 vs. 2), SEL1L ΔAPH failed to reduce aggregates, similar to that of SEL1L L709P (lane 4 and 7 vs. 2, Fig. S4E). Collectively, these results demonstrate that SEL1L L709P abolishes the SEL1L-HRD1 interaction via the disruption of APH.

### SEL1L L709P abolishes the SEL1L-HRD1 interaction in human cells

To confirm these findings in human cells, we generated biallelic *SEL1L*^*L709P*^ KI HEK293T cells using CRISPR/Cas9 (Fig. S5A-B). SEL1L and HRD1 KO HEK293T cells were included as controls (26). Consistent with the mouse findings, SEL1L protein levels increased by ∼30-40%, while HRD1 levels were decreased ∼40%, and OS9 levels increased fivefold (Fig. S5C, quantified in Fig. S5D). Known ERAD substrates, including IRE1α and CD147, were elevated by ∼50%, phenocopying *SEL1L* or *HRD1* KO cells (Fig. S5C, quantified in Fig. S5D). Immunoprecipitation of endogenous SEL1L confirmed the loss of HRD1 binding in *SEL1L*^*L709P*^ KI cells, with OS9 and UBE2J1 interactions largely intact (Fig. S5E, quantified in Fig. S5F). Together, these data demonstrate that the L709P mutation selectively disrupts SEL1L-HRD1 interaction in both mouse and human cells.

### The SEL1L-HRD1 interaction is required for substrate and E2 enzyme recruitment

To further assess functional consequence of SEL1L L709P mutation, we examined the ERAD of proAVP-G57S-HA in WT, KI and KO HEK293T cells. Immunoprecipitation of proAVP revealed that it associated with OS9 and SEL1L, but not HRD1, in *SEL1L*^*L709P*^ KI HEK293T cells (lanes 3-5, Fig. 6A and quantified in Fig. 6B), indicating impaired substrate transfer to HRD1. HRD1 immunoprecipitation revealed that SEL1L L709P disrupted not only SEL1L binding, but also recruitment of E2 ubiquitin-conjugating enzyme UBE2J1 (Fig. 6C). As a result, proAVP polyubiquitination was severely impaired, mirroring defects in *SEL1L*^*-/-*^ and *HRD1*^*-/-*^ cells (Fig. 6D). Together, these results demonstrate that the SEL1L- HRD1 interaction is required for both substrate and E2 enzyme recruitment.

**Figure 6.**
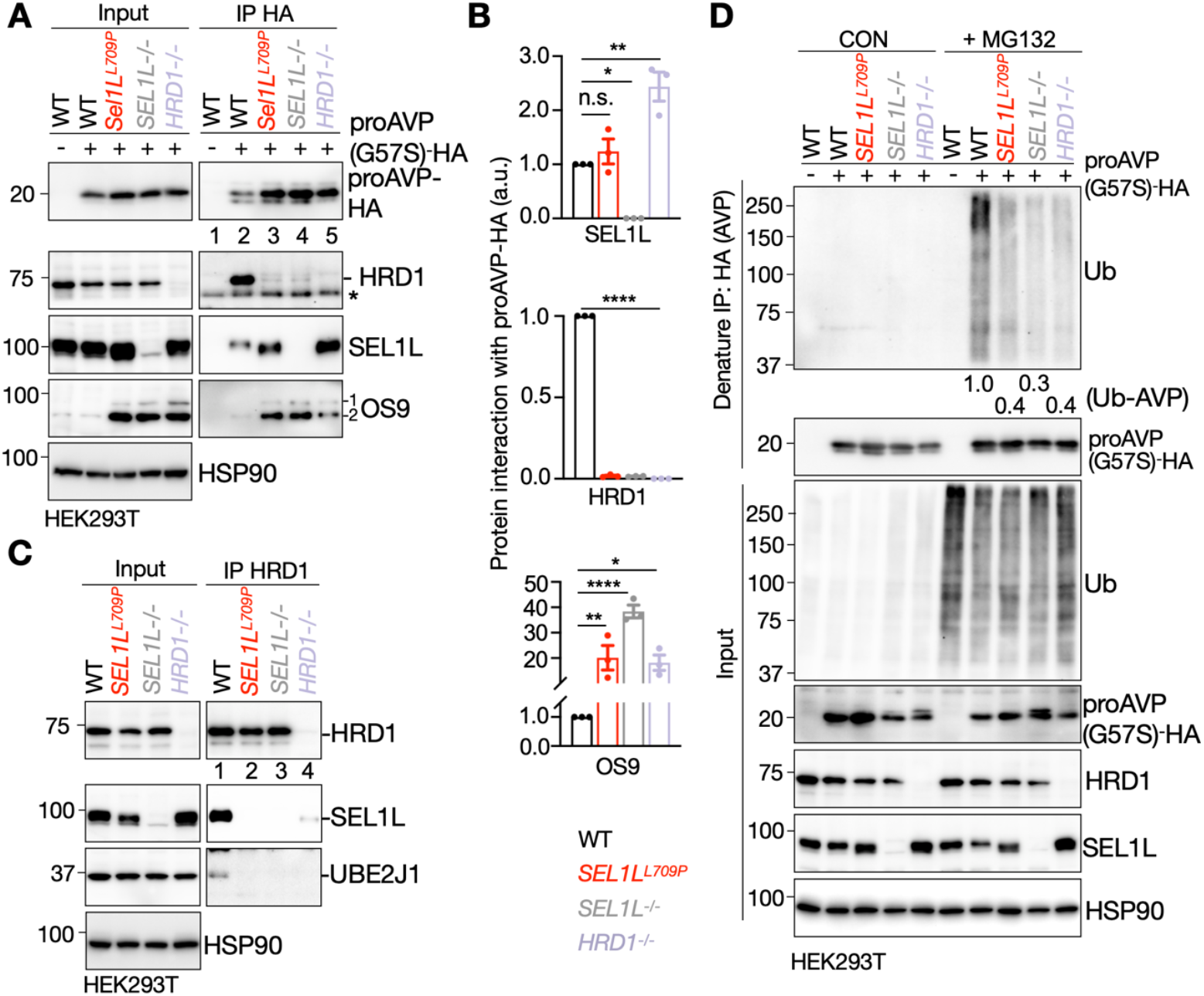
The SEL1L-HRD1 interaction is essential for substrate recruitment and ubiquitination. (A-B) Co-immunoprecipitation of proAVP (G57S)-HA in HEK293T cells showing the interactions with OS9 and SEL1L across all lines, but not HRD1 in *SEL1L*^*–/–*^ and *SEL1L*^*L709P*^ KI HEK293T cells (A). Quantification is shown in (B). HSP90 serves as the input loading control. Data represents n = 3 independent experiments, with 2 repeats. (C) Co-immunoprecipitation of HRD1 in *HRD1*^*–/–*^, *SEL1L*^*–/–*^ and *SEL1L*^*L709P*^ KI HEK293T cells showing the loss of its interaction with UBE2J1. HSP90 serves as the input loading control. Data represents n = 3 independent experiments, with 2 repeats. (D) Immunoblot of ubiquitinated proAVP (G57S)-HA in *HRD1*^*–/–*^, *SEL1L*^*–/–*^ and *SEL1L*^*L709P*^ KI HEK293T cells treated with or without 10 μM MG132 for 4 hours. The quantification of AVP ubiquitination shown below the gel and represents the average of n = 3 independent experiments, with 2 repeats. HSP90 serves as the input loading control. Values, mean ± SEM. n.s., not significant; ^*^p < 0.05; ^**^p < 0.01; ^***^p < 0.001; ^****^p < 0.0001 using one-way ANOVA with Dunnett’s multiple comparisons test.

### HRD1 overexpression partially rescues ERAD defects in *SEL1L*^*L709P*^ cells

Inspired by yeast studies in which Hrd1 overexpression bypasses loss of Hrd3 (23, 29), we tested whether increased HRD1 expression can rescue ERAD defects in mammalian cells. Overexpression of HRD1 in *SEL1L*^*L709P*^ KI and *SEL1L*^*-/-*^ HEK293T cells restored the degradation of an endogenous ERAD substrate CD147 (45) in a dose-dependent manner (Fig. 7A and quantified in Fig. 7B). Cycloheximide chase confirmed that HRD1 overexpression accelerated CD147 turnover (Fig. S6A-B), and polyubiquitination of CD147 was similarly enhanced (Fig. S6C) in both *SEL1L*^*L709P*^ and *SEL1L*^*–/–*^ HEK293T cells.

**Figure 7.**
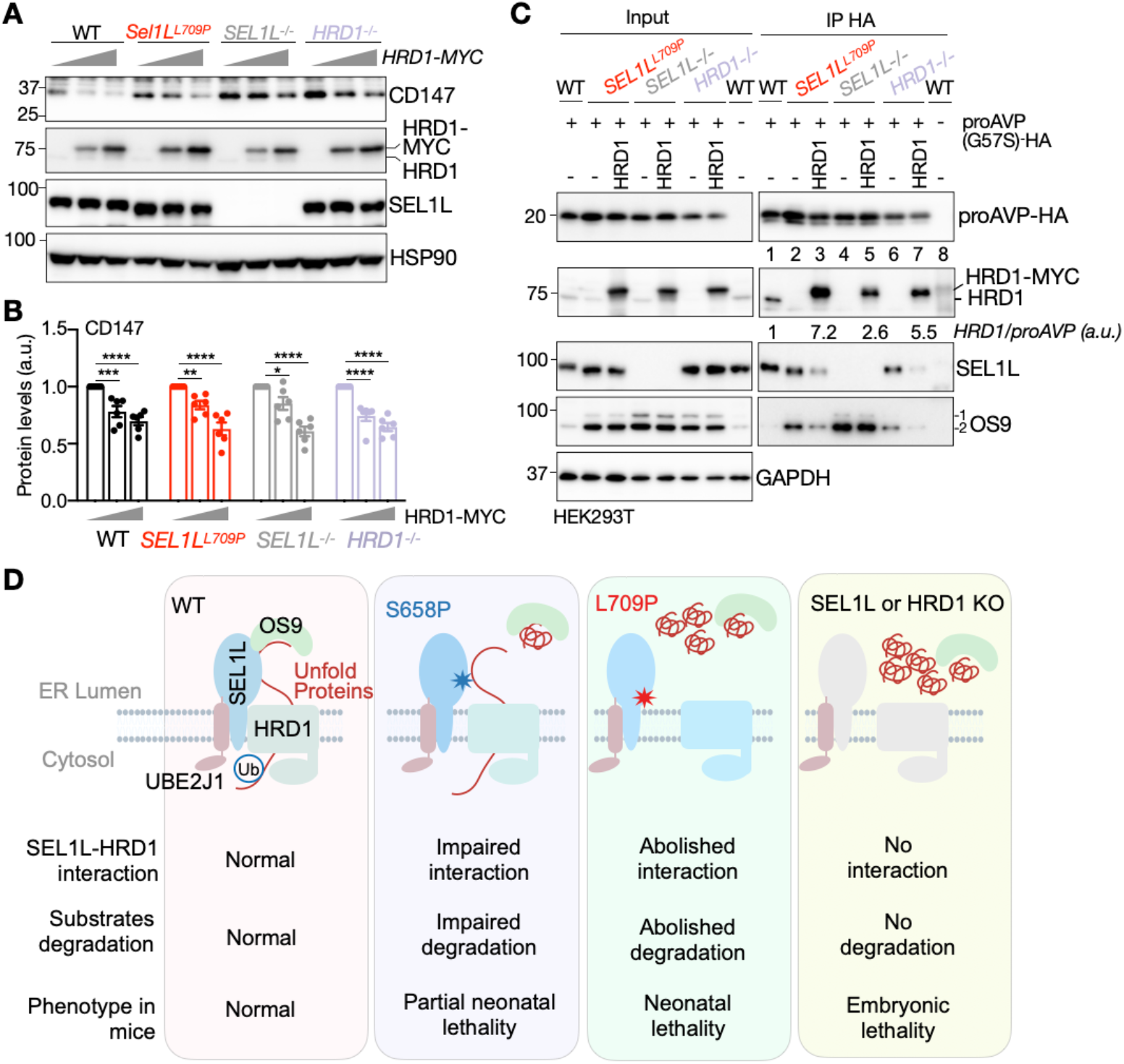
HRD1 overexpression fails to rescue ERAD deficiency in SEL1L L709P cells and the proposed model. (A–B) Western blot analysis of CD147, an endogenous ERAD substrate, in WT, SEL1L^−/−^, SEL1L L709P, and HRD1^−/−^ HEK293T cells with increasing amounts of wild-type HRD1 (A). Quantification is shown in (B), normalized to the loading control HSP90; Data represents n = 6 independent experiments, with 2 repeats. (C) Co-IP of proAVP(G57S)-HA with HRD1, with or without HRD1-MYC overexpression, in WT, SEL1L^-/-^, SEL1L^L709P^, and HRD1^-/-^ cells. Quantification of HRD1 shown below the gel represents the average of n = 3 independent experiments, normalized to proAVP. GAPDH serves as the input loading control. (D) Model illustrating the correlation between SEL1L-HRD1 ERAD activity and perinatal lethality in mice. Values are presented as mean ± SEM. n.s., not significant; ^*^p < 0.05; ^**^p < 0.01; ^***^p < 0.001; ^****^p < 0.0001 using two-way ANOVA with Tukey’s post hoc test (B).

Lastly, we performed co-immunoprecipitation assays to assess HRD1–substrate interactions in proAVP-HA-transfected *SEL1L*^*L709P*^ KI and *SEL1L*^*–/–*^ HEK293T cells with or without HRD1 overexpression. Immunoprecipitation of proAVP revealed that while its association with endogenous HRD1 was abolished in both SEL1L L709P KI and *SEL1L*^*-/-*^ cells (lanes 2 and 4 vs. 1), it was restored upon HRD1 overexpression (lanes 3 vs. 2 and 5 vs. 4, Fig. 7C). Together, these results demonstrate that while SEL1L is essential for efficient HRD1-mediated ERAD under physiological conditions, supraphysiological HRD1 levels can partially bypass the requirement for SEL1L in substrate processing in mammalian cells.

## DISCUSSION

Despite extensive *in vivo* evidence highlighting the importance of SEL1L and HRD1, definitive proof that SEL1L is required for HRD1 function in mammals has remained elusive. This gap stems largely from the absence of models allowing selective disruption of the SEL1L–HRD1 interaction without globally affecting the abundance of either protein. In this study, we characterize the disease-associated SEL1L variant L709P, which abolishes HRD1 binding while preserving SEL1L protein levels and maintaining HRD1 expression at ∼50% of normal levels. This approach allowed us to isolate the functional significance of the SEL1L–HRD1 interface in mammalian ERAD and physiology.

Our first major finding is that SEL1L is essential for HRD1 function in mammals (Fig. 7D). Specifically, we show that: (a) SEL1L is indispensable for HRD1-mediated ERAD under physiological conditions; and (b) the extend of SEL1L-HRD1 disruption directly correlates with the severity of ERAD impairment. Mechanistically, SEL1L is required for recruiting both OS9-bound substrates and the E2 enzyme UBE2J1 to HRD1. Disruption of this interaction impairs substrate engagement and efficient ubiquitin transfer, leading to ERAD failure and perinatal lethality in mice. Importantly, supraphysiological expression of HRD1 partially rescues these defects – mirroring similar compensatory mechanisms observed in yeast (23, 29). However, unlike in yeast where deletion of *hrd3* does not abolish substrate association with Hrd1 (24), SEL1L is strictly required for HRD1-substrate interaction in mammals, highlighting an evolutionary divergence in ERAD architecture. Our findings support a model in which SEL1L functions as a scaffolding hub in higher eukaryotes, coordinating substrate recognition and ubiquitin conjugation within the ERAD complex. Nonetheless, given the cellular diversity in mammals, it is plausible that this requirement may be bypassed in certain contexts through unknown mechanisms or cell-type-specific adaptations.

Our second major finding links the degree of ERAD impairment to organismal viability, supporting a graded, dose-dependent model of ERAD dysfunction (Fig. 7D). Germline deletion of either *Sel1l* or *Hrd1* results in embryonic lethality, whereas acute deletion in adults leads to death within weeks (13-16). Previously, we reported that the hypomorphic *Sel1l* S658P variant causes partial neonatal lethality and early-onset neurodegeneration in survivors (26). Here, we show that the L709P variant completely abolishes the SEL1L- HRD1 interaction and causes a more severe ERAD defect than S658P. This molecular disruption is mirrored *in vivo*: L709P homozygosity results in complete neonatal lethality, unlike the partial viable S658P KI mice (Fig. 7D). These observations underscore the functional necessity of the SEL1L-HRD1 interaction and reinforce the importance of ERAD integrity for postnatal survival. Notably, this dose-dependent relationship is consistent with clinical observations in ERAD-associated neurodevelopmental disorders with onset in infancy (ENDI) patients, where disease severity inversely correlates with residual ERAD activity in individuals harboring biallelic hypomorphic variants in *SEL1L* or *HRD1* (20-22).

The precise cause of neonatal lethality in *SEL1L*^*L709P*^ KI mice remains unclear. Histological examination of major tissues appears largely normal, and ultrastructural analysis reveals only mild ER dilation. Despite clear defects in SEL1L-HRD1 ERAD, activation of the UPR is subtle, consistent with previous reports showing that SEL1L-HRD1 ERAD dysfunction does not robustly trigger an overt UPR signaling (17-19, 22). This attenuated response may reflect compensatory mechanisms, such as chaperone upregulation (46) or engagement of alternative degradation pathways such as autophagy (19, 40, 41). As neonatal survival depends on critical physiological functions such as respiration, thermoregulation, and suckling (47), further studies are needed to determine whether failure in SEL1L-HRD1 ERAD directly compromises these functions.

In summary, our work provides direct evidence that the SEL1L-HRD1 interaction is essential for mammalian ERAD and postnatal viability. These findings resolve a long-standing question in ERAD biology and uncover a structurally defined node that may serve as a therapeutic target for diseases associated with SEL1L-HRD1 ERAD dysfunction and ER proteostasis dysfunction.

## MATERIALS AND METHODS

### Mouse Models

SEL1L S658P KI mice were previously described (26) and hepatocyte-specific Sel1L KO mice were previously described (35-37). KI mouse models harboring SEL1L P695T and L705P mutations (corresponding to human P699T and L709P, respectively) were generated on the B6/SJL background via CRISPR/Cas9 at the University of Michigan Molecular Genetics Core. Two single-guide RNAs (sgRNAs) targeting exon 20 of the Sel1L gene were designed using CRISPOR (http://crispor.tefor.net) (sgRNA1: <u>CTAGGACATTCACCTTGCAA; sgRNA2: AAGCAAACGTAAGTGAGCCG</u>), synthesized via the Synthego sgRNA Synthesis Kit, and validated in fertilized mouse oocytes. The donor template (Ultramer dsDNA, IDT) contained the desired mutation and silent mutations to facilitate homology-directed repair:

Donor 1 (P695T):

ACACAAATTTTTCTGAAACATTTCCTTTGCTCTCTGTCCTCTAGGATATTCATCTTGC TAAACGCTTTTATGACATGGCAGCCGAAGCTAGC**ACA**GATGCACAAGTACCTGTGT TCCTCGCACTCTGCAAATTAGGTGTCGTCTATTTCTTACAGTACATACGGGAGGCCAATGTAAGTGAGCCGTTGCTCTTAGTTACA;

Donor 2 (L705P):

ACACAAATTTTTCTGAAACATTTCCTTTGCTCTCTGTCCTCTAGGATATTCATCTTGCTAAACGCTTTTATGACATGGCAGCCGAAGCTAGCCCAGATGCACAAGTACCTGTGT TCCTCGCA**CCC**TGCAAATTAGGTGTCGTCTATTTCTTACAGTACATACGGGAGGCC

AATGTAAGTGAGCCGTTGCTCTTAGTTACA

This template was co-injected with recombinant Cas9 protein (Sigma-Aldrich) and sgRNAs into B6/SJL oocytes, followed by embryo transfer into pseudopregnant females. Founder mice were genotyped by Sanger sequencing (F: 5′- TTAGGTCGGATCTTGAAAGGCTAAC-3′; R: 5′-CACAGGTTACCCACGAGGAATTC-3′) and backcrossed to C57BL/6J for three generations prior to generating homozygous KI lines and littermate controls.

All mouse strains used in this study were on a C57BL/6J genetic background. Mice were housed in a specific pathogen-free facility at 22 ± 1 °C under a 12-hour light/dark cycle with 40–60% humidity and fed a low-fat diet (13% fat, 57% carbohydrate, 30% protein; LabDiet 5LOD). All animal procedures were approved by the University of Virginia (Protocol #4459) and were conducted in accordance with the guidelines of the National Institutes of Health (NIH).

### Neonatal Survival Curve Analysis

To assess neonatal survival, pups were monitored from birth (postnatal day 0, P0) through the first 48 hours of life. Litters were inspected at every three hours, and the number of surviving pups was recorded at each designated time point. Dead pups were removed at each inspection, and tail biopsies were collected immediately for genotyping. After 48 hours, tail samples were collected from all surviving pups. Pups exhibiting spontaneous movement and active breathing were considered alive. Genotyping was performed on tail biopsies collected from both surviving and non-surviving pups. Kaplan–Meier survival curves were generated using GraphPad Prism software, with survival plotted as the percentage of live pups over time. Statistical significance between genotypes was determined using the log-rank (Mantel–Cox) test.

### Histological Analysis

P0 mice were fixed following a modified protocol based on the Mouse Phenotyping Core at UCSD (https://mousepheno.ucsd.edu/histo.shtml). Briefly, mice were subjected to a small abdominal incision and perfused with 15 mL of freshly prepared fixative consisting of absolute ethanol, 37% formaldehyde, and glacial acetic acid in a 6:3:1 ratio. After perfusion, the mice were incubated in the same fixative overnight at 4 °C. Tissues were then post-fixed in freshly prepared fixative for an additional 3 days, with daily replacement of the solution. To enhance fixation quality, tissues were sagittally sectioned and further fixed in 10% neutral-buffered formalin (Sigma) for an additional 3 days prior to processing. Following fixation, tissues were embedded in paraffin, sectioned, and stained with hematoxylin and eosin (H&E) at the UVA Research Histology Core.

### Generation of MEFs

MEFs were generated from embryonic day 15.5 (E15.5; counting noon of the day when the vaginal plug was found as E0.5) embryos obtained from matings between heterozygous mice carrying different SEL1L variants (P699T, S658P, or L709P) as previously published (16). Briefly, the head and liver were removed from each embryo (and retained for genotyping), and the remaining tissue was finely minced and digested in 5 mL of trypsin at 37°C for 10 minutes. The digested tissue was then pipetted vigorously to obtain a single-cell suspension and plated in 10-cm culture dishes. MEFs were cultured in DMEM (Gibco) supplemented with 10% fetal bovine serum (FBS; Fisher Scientific) at 37°C in a humidified incubator with 5% CO₂, and expanded for experiments. Inducible Sel1L-deficient MEFs (*Sel1L*^*ERCre*^) were generated as previously described (16).

### CRISPR/Cas9-mediated Gene Knockout and KI

HEK293T cells (ATCC) were cultured in DMEM (Gibco) supplemented with 10% fetal bovine serum (FBS; Fisher Scientific) at 37°C in a humidified incubator with 5% CO₂. SEL1L and HRD1 knockout HEK293T cell lines were generated following previously published methods(26). Guide RNAs were cloned into lentiCRISPR v2 (Addgene #52961) and transfected using polyethyleneimine (PEI, 5 μL/μg DNA). After puromycin selection (2 μg/mL for 24 h), cells were expanded for downstream analyses. Guide RNAs:

SEL1L,

F: 5′-CACCGGGCTGAACAGGGCTATGAAG-3′;

R: 5′-AAAC CTTCATAGCCCTGTTCAGCCC-3′

HRD1,

F: 5′-CACCGGGCCAGCCTGGCGCTGACCG-3′;

R: 5′-AAACCGGTCAGCGCCAGGCTGGCCC-3′

SEL1L L709P KI HEK293T cells were generated via CRISPR-Cas9-Homology-Directed Repair (CIRSPR-Cas9-HDR) as we previously described (26). crRNA (guide sequence): 5’- GACGACGCCCAATTTGCAGA -3’

HDR Donor Oligo (the desired mutation is underlined): 5’- TCCCGTATGTACTGCAAGAAATAGACGACGCCCAATTTGCAGGGGGCTAGGAAGACTGGAACTTGTGCATCTGGGCTGGCTTC-3’

Amplification PCR primers:

F: 5’- AGCTTGGCATTTTTGTTTAGGTG -3’

R: 5’- CGTCAGGAAGTGTCAAACGCT -3’

Sequencing primer:

5’- TTTGACAAGAAATGCATTTTTG -3’

### Phos-tag SDS-PAGE and Western blot

Phosphorylated IRE1α was resolved using Phos-tag SDS-PAGE (5% gel containing 50 μM Phos-tag and 50 μM MnCl_₂_) as previously described (48, 49). Gels were treated with 1 mM EDTA prior to transfer.

### Drug Treatment

MEFs or HEK293T cells were treated with 50 μg/mL cycloheximide or 10 μM MG132 for the indicated time points, followed by Western blot or immunoprecipitation.

### Immunoprecipitation

MEF and liver lysates were prepared in lysis buffer (150 mM NaCl, 25 mM Tris-HCl pH 7.5, 0.2% NP-40, 0.1% Triton X-100), incubated with anti-SEL1L or anti-HRD1 antibodies overnight at 4°C, followed by Protein A agarose (Invitrogen). Bound proteins were washed three times with lysis buffer (5 min each on a rotator at 4°C), then eluted using 2× SDS sample buffer (prepared by diluting 5× SDS sample buffer with lysis buffer), and subsequently analyzed by SDS-PAGE and Western blotting. For ubiquitination assays, lysates from transfected cells expressing proAVP-G57S-HA were prepared in denaturing buffer (150 mM NaCl, 50 mM Tris-HCl pH 7.5, 1 mM EDTA, 1% NP-40, 1% SDS, 5 mM DTT, protease inhibitors), diluted 10-fold in the former lysis buffer, and incubated with anti-HA agarose (Sigma A2095) overnight at 4°C. Bound proteins were washed and eluted using the same conditions as described in the previous immunoprecipitation experiment, and subsequently analyzed by SDS-PAGE and Western blotting. The intensity of the co-immunoprecipitated (prey) band was normalized to the bait signal to quantify the binding efficiency.

### Non-reducing SDS-PAGE

Cell lysates were prepared in NP-40 buffer (50 mM Tris-HCl pH 8.0, 0.5% NP-40, 150 mM NaCl, 5 mM MgCl2 and protease inhibitor) supplemented with 10 mM N-ethylmaleimide, incubated at 37°C for 30 min, and resolved on 6–15% gradient gels under non-reducing conditions as previously described (37).

### TEM

Mice were perfused with 4% glutaraldehyde and 4% paraformaldehyde in 0.1 M cacodylate buffer. Tissues were post-fixed overnight in 3% glutaraldehyde and 3% paraformaldehyde in 0.1 M Sorenson’s buffer. Following fixation, tissues were embedded, sectioned, and stained with uranyl acetate and lead citrate by the University of Virginia Molecular Electron Microscopy Core. Images were acquired using a Tecnai F20 transmission electron microscope at the same facility.

## Statistical Analysis

All statistical analyses were performed using GraphPad Prism 8.0. Results are presented as mean ± SEM. Statistical significance was determined using two-tailed Student’s t-test (for two-group comparisons) or one-/two-way ANOVA with post hoc Dunnett’s or Tukey’s multiple comparisons test, as appropriate. A p-value < 0.05 was considered statistically significant.

## ACKNOWLEDGMENTS

We thank Dr. David Castle, and members of the Qi, Sun, and Arvan laboratories for insightful discussions; Dr. Zhengfeng Song and Shengyi Sun for sharing the mice; Drs. Michael Purdy and David Cooper for assisting with the F20 training and data collection; Dr. Sara Wilmsen for TEM sample preparation; and University of Virginia School of Medicine Research Histology Core Facility and Molecular Electron Microscopy Core Facility (NIH grant G20-RR31199) for sample processing and data collection. The work is supported by 1R01AG089640 and 1R01NS138119 (L.Q.). H.H.W. is supported by Alzheimer’s Association Postdoctoral Research Fellowship (25AARF-1375486).

