## Supplemental for "Functional integrity of the SEL1L-HRD1 complex is critical for ERAD and organismal viability"

**This PDF file includes:**

Supporting text
SI References
Tables S1
Figures S1 to S6

### Supporting Information Text

#### Methods

##### Genetic Analysis

Evolutionary conservation of these residues was assessed using Clustal W and visualized with Jalview (1). Orthologous protein sequences from human (NP_005056.3), chimpanzee (JAA44458.1), pig (XP_020955243), mouse (NP_001034178.1), Drosophila (NP_001262882.1), yeast (QHB10358.1), and chicken (XP_040558090.1) were aligned.

##### Mouse Genotyping

Genomic DNA from tail biopsies was amplified using the following primers:
WT-F 5′-CTGTCCTCTAGGACATTCACCTTGCA-3′;
WT-R 5′-CAGAAACCACCTGCACCGAT-3′;
KI-F 5′-CTGTCCTCTAGGATATTCATCTTGCT-3′;
KI-R 5′-CAGAAACCACCTGCACCGAT-3′).
L709P and P699T KI mice shared the same primer sets for genotyping. Genotyping procedure and primers for SEL1L S658P KI mice were previously described (2).

##### Clinical Data and Analysis

Human SEL1L variants (P699T and L709P) were identified through the Baylor College of Medicine's clinical genomic sequencing program. Position-specific scoring matrix (PSSM) scores were calculated using PSI-BLAST, and pathogenicity predictions were generated using PolyPhen-2 (3).

##### Plasmids and Transfection

Mouse Sel1L WT and S658P in the pcDNA3 expression vector with a C-terminal FLAG tag were previously reported (2). Point mutations (P699T, L709P, and S658P) and deletion mutation ΔAPH (Δ699-720) of Sel1L were introduced by site-directed mutagenesis (2) and verified by Sanger sequencing. Primers to generate the mutants are:
P699T: F 5′-GGCAGCCGAAGCTAGCACAGATGCACAAG-3′,
R 5′-CTTGTGCATCTGTGCTAGCTTCGGCTGCC-3′)

L709P: F 5′-CCTGTGTTCCTCGCACCCTGCAAATTAGGTG-3′,
R 5′-CACCTAATTTGCAGGGTGCGAGGAACACAGG-3′,

ΔAPH: F 5′-ACATGGCAGCCGAAGCTAGCATACGGGAAGCAAACATTCGA,
R 5′-GCTAGCTTCGGCTGCCATGT

For most experiments, HEK293T cells were seeded at a density of 1 × $10^6$ cells per well in six-well plates or 3 × $10^6$ cells per 6-cm dish. After 24 hours, the medium was replaced with fresh, pre-warmed DMEM, and cells were transfected using PEI at a ratio of 5 μL PEI per 1 μg plasmid DNA. Cells were harvested 24 hours later or subjected to drug treatments as indicated. For the transfection of SEL1L variants (*SEL1L-FLAG*, *SEL1L-L709P-FLAG*, *SEL1L-P699T-FLAG*, *SEL1L-S658P-FLAG*, and *SEL1L-ΔAPH-FLAG*), 1 μg of plasmid DNA was used per well (six-well plate) or 3 μg per 6-cm dish. For proAVP-G57S-HA plasmid, 1 μg of plasmid DNA was used per well (six-well plate) or 3 µg per 6-cm dish. For transfection of the human HRD1-MYC plasmid, 0.2 µg and 0.5 µg per well (six-well plate) were used for Fig. 7A, 0.5 µg per well was used for Fig. S6A, and 2 µg per 6-cm dish was used for Fig. 7C and Fig. S6C.

**Immunoblotting**

Mouse tissues and MEFs were lysed in Triton X-100 buffer (50 mM Tris-HCl pH 7.5, 150 mM NaCl, 1% Triton X-100, 1 mM EDTA) supplemented with protease/phosphatase inhibitors (Sigma). Lysates were cleared by centrifugation (16,000 g, 10 min), and protein concentrations of supernatants were measured using Bio-Rad Protein Assay Dye (Bio-Rad). A total of 20–50 µg of protein was denatured at 95°C for 5 minutes in 5x SDS sample buffer containing 250 mM Tris-HCl (pH 6.8), 10% SDS, 0.05% bromophenol blue, 50% glycerol, and 1.44 M β-mercaptoethanol. SDS-PAGE and Western blotting were performed using standard protocols (4, 5). The blots were incubated in 2% BSA/Tri-buffered saline tween-20 (TBST) with primary antibodies overnight at 4°C: anti-HSP90 (Santa Cruz, #sc-13119, 1:5,000), anti-GAPDH (Proteintech, #60004-1, 1:5000), anti-SEL1L (home-made, 1:10,000) (6), anti-HRD1 (Proteintech, #13473-1, 1:2,000), anti-OS9 (Abcam, #ab109510, 1:5,000), anti-CD147 (Proteintech, #11989-1, 1:3,000), anti-IRE1α (Cell Signaling, #3294, 1:2,000), anti-UBE2J1 (Santa Cruz, #sc-377002, 1:3,000), anti-HA (Sigma, #i2018, 1:1,000), anti-PERK (Cell Signaling, #3192, 1:5000), anti-eIF2α (Cell Signaling, #9722, 1:5000), anti-p-eIF2α (Cell Signaling, #9721, 1:1000) and anti-ubiquitin (Santa Cruz, #sc-8017, 1:1000). Membranes were washed with TBST and incubated with the appropriate HRP-conjugated secondary antibodies at room temperature for 1 hour prior to detection using an ECL chemiluminescence system (Bio-Rad). For general Western blot, HRP-conjugated secondary antibodies (Bio-Rad, 1:10,000) were used. For immunoprecipitation samples, either anti-Rabbit IgG TrueBlot HRP (Rockland, #18-8816-33, 1:500) or anti-Mouse IgG TrueBlot HRP (Rockland, #18-8817-31, 1:500) was used as appropriate. Band intensity was quantified using Image Lab software (Bio-Rad).

Protein lysates from the livers of 8-week-old mice treated with tunicamycin (TM, 1 mg/kg, i.p.) for 24 hours as a positive control for UPR. For phosphatase treatment, 100 µg of tissue lysate was incubated with 1 µl of lambda phosphatase (λPPase; New England BioLabs, catalog #P0753S) in 1× PMP buffer (New England BioLabs, catalog #B0761S) supplemented with 1 mM $MnCl_2$ (New England BioLabs, catalog #B1761S) at 30 °C for 60 minutes. The reaction was terminated by adding 5× SDS sample buffer and heating at 90 °C for 5 minutes, as previously described (7).

**RNA Extraction, RT-PCR and Q-PCR**

Total RNA was extracted using TRI Reagent (Sigma) as previously described (2). *Xbp1* mRNA splicing was analyzed by RT-PCR using intron-flanking primers (F: 5′-ACGAGGTTCCAGAGGTGGAG-3′; R: 5′-AAGAGGCAACAGTGTCAGAG-3′) as previously described (8). PCR products were resolved by agarose gel electrophoresis and quantified with Image Lab. For qPCR, gene expression levels were normalized to *L32*. Primer sequences were as follows:

*L32* F: 5’-GAGCAACAAGAAAACCAAGCA-3’; R: 5’-TGCACACAAGCCATCTACTCA-3’.
*Sel1L* F: 5’-TGGGTTTTCTCTCTCTCCTCTG-3’; R: 5’-CCTTTGTTCCGGTTACTTCTTG-3’.
*OS9* F: 5’-GCTGGCTGACTGATGAGGAT-3’; R: 5’-CGGTAGTTGCTCTCCAGCTC-3’.

*Hrd1* F: 5’-AGCTACTTCAGTGAACCCCACT-3’; R: 5’-CTCCTCTACAATGCCCACTGAC-3’.
*Hrd1* F: 5’-5ATCTGCGCAAATTCAGAACC-3’; R: 5’-CTCCATGGCTTGGTAGGTGT-3’.
*L32* F: 5’-GAGCAACAAGAAAACCAAGCA-3’; R: 5’-TGCACACAAGCCATCTACTCA-3’.

### Tables

**Table S1. Clinical characteristics of patients harboring the SEL1L L709P or P699T variant.**

| | Patient 1 | Patient 2 |
|---|---|---|
| Sex | F | M |
| Age (month) | 0.836 | 3.599 |
| De-ID phenotype | Intracranial hemorrhage, retinal hemorrhage, recurrent fractures, blue sclerae, and premature birth. | Autism spectrum disorder, history of speech delay, macrocephaly, hypotonia, abnormal MRI suggestive of hemimegalencephaly. Family history is significant for mother with branchio-oto-renal syndrome and hearing loss. |
| Variant type | Novel | Novel |
| Gene | SEL1L | SEL1L |
| Nucleotide | c.2095C>A | c.2126T>C |
| Amino acid change | p.P699T | p.L709P |
| Zygosity | Heterozygous | Heterozygous |
| Comments | Novel variant | Novel variant. Mother is negative. Father is heterozygous. |
| PolyPhen-2 | Damaging | Damaging/Probably damaging |

### Supplemental Figures

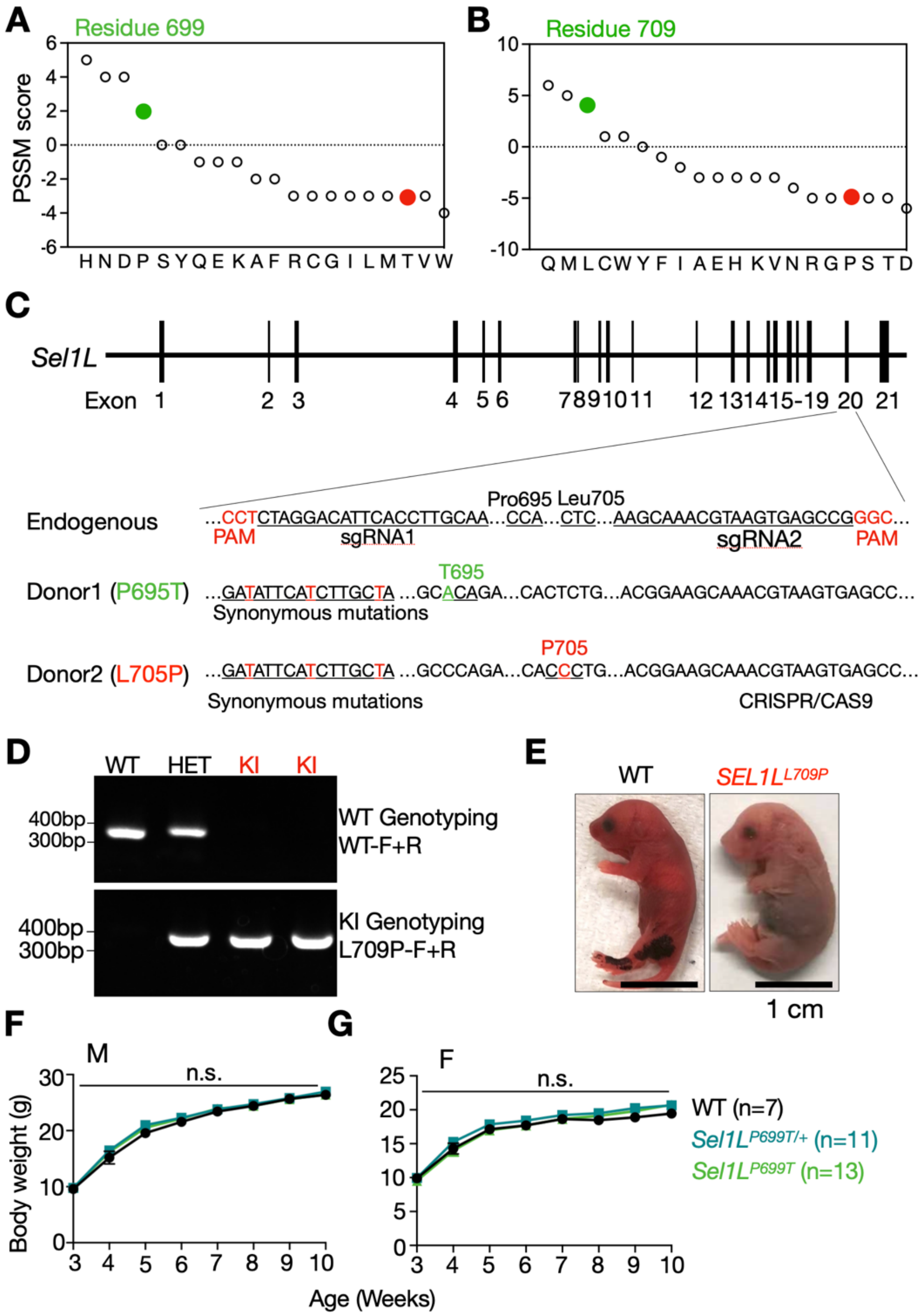


**Figure S1. Generation and characterization of *Sel1L*$^{L709P}$ and *Sel1L*$^{P699T}$ KI mice.**
(A–B) Position-specific scoring matrix (PSSM) analysis of SEL1L residues P699 (A) and L709 (B) (green) and variants (red).
(C) Schematic diagram showing CRISPR/Cas9-mediated gene editing. Mouse residues P695 and L705 correspond to human P699 and L709, respectively.
(D) Representative genotyping PCR showing the identification of WT, heterozygous (HET), and homozygous KI alleles. L709P and P699T KI mice shared the same primer pairs for genotyping.
(E) Gross examination of deceased *Sel1L*$^{L709P}$ pups reveals no external malformations.
(F-G) Growth curves of *Sel1L*$^{P699T}$ male (F) and female (G) mice. Values are presented as mean ± SEM. n.s., not significant, using two-way ANOVA with Tukey's post hoc test.

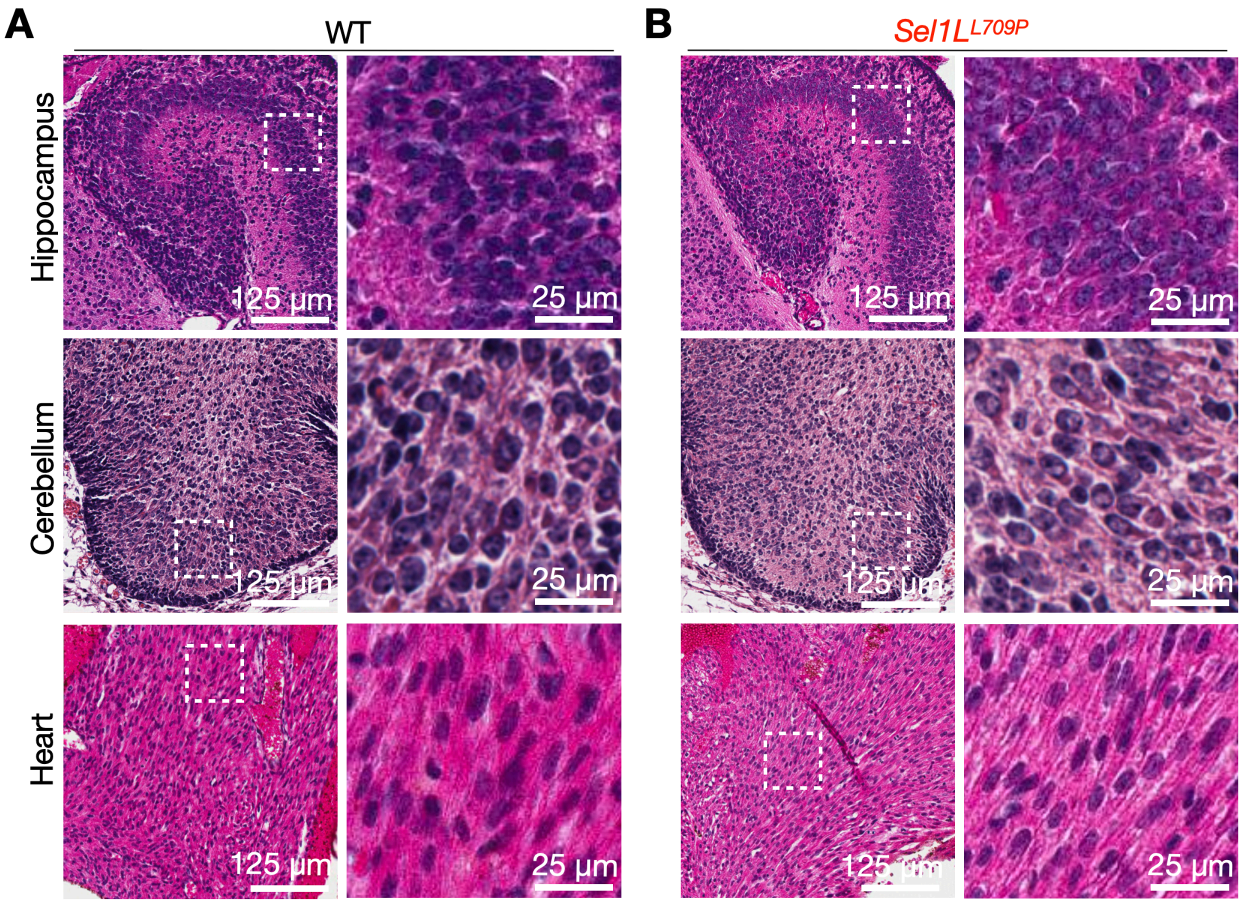


**Figure S2. Histological examination of WT (A) and *Sel1L$^{L709P}$* KI (B) neonatal tissues including hippocampus, cerebellum and heart.** Representative images from three independent P0 mice per genotype with no overt abnormalities observed.

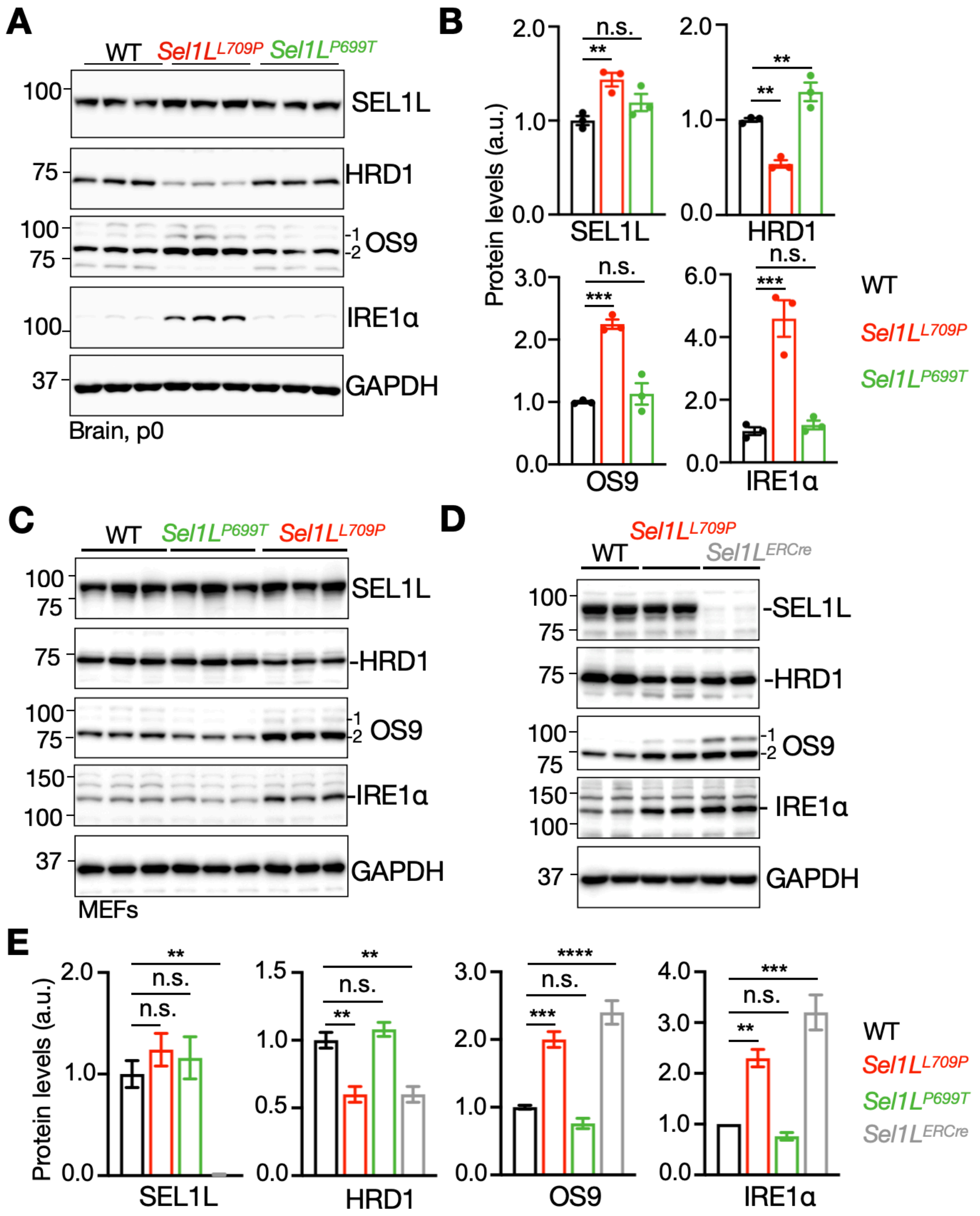


**Figure S3. Impaired ERAD function in *Sel1L$^{L709P}$* KI MEFs and brain tissue.**

(A-B) Immunoblot analysis of ERAD-related proteins in whole-brain lysates from WT and two KI mice (A). Quantification is shown in (B), normalized to the loading control GAPDH. N = 3 mice per group, with 2 repeats.

(C-E) Immunoblot analysis of ERAD components and substrates in WT, *Sel1L$^{L709P}$* and *Sel1L$^{P699T}$* and *Sel1L$^{ERCre}$* MEFs (C-D). Quantification is shown in (E), normalized to GAPDH. N = 3 sets of MEFs per genotype, with 2 repeats.

Values are shown as mean ± SEM. n.s., not significant; *$p < 0.05$; **$p < 0.01$; ***$p < 0.001$; ****$p < 0.0001$ using one-way ANOVA with Dunnett's multiple comparisons test.

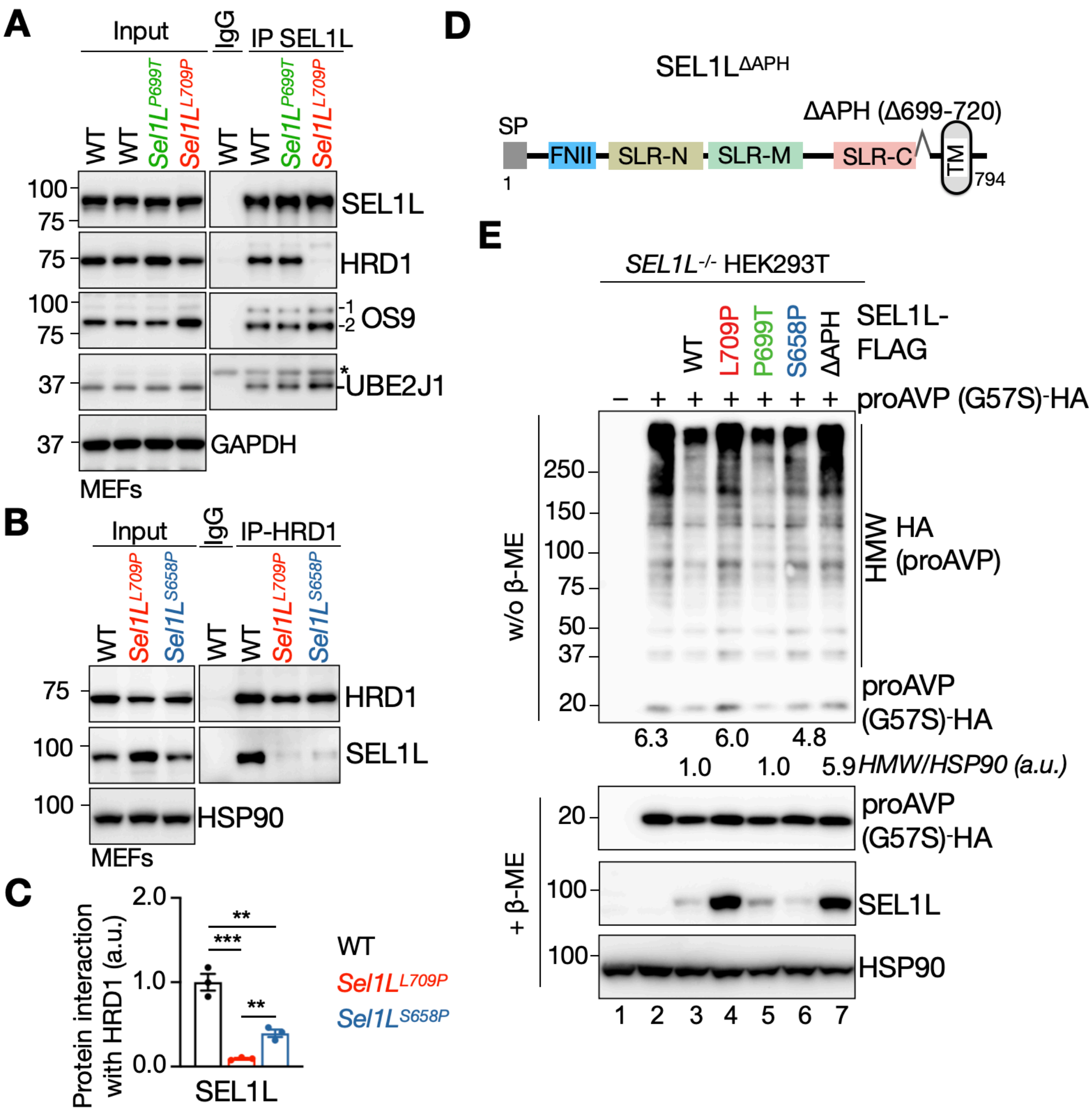


**Figure S4. SEL1L L709P mutation abolishes SEL1L-HRD1 interaction *in vitro*.**
(A) Co-IP of endogenous SEL1L from MEFs of WT, *Sel1L*$^{P699T}$, and *Sel1L*$^{L709P}$ mice, followed by immunoblotting for ERAD components. Quantification is shown in Fig. 5E, normalized to GAPDH. Non-specific bands are indicated by asterisks.
(B-C) Co-IP of endogenous HRD1 from MEFs of the indicated genotypes, analyzed for interaction with SEL1L (B). Quantification is shown in (C), normalized to HRD1 levels; Data represents the MEFs from n = 3 mice per group, with 2 repeats. HSP90 is the loading control.
(D) Schematic diagram of the human SEL1L ΔAPH protein domains.
(E) Reducing and non-reducing SDS-PAGE followed by immunoblotting to detect HMW aggregates of the misfolded ERAD substrate proAVP-G57S in *SEL1L*$^{-/-}$ HEK293T cells expressing the indicated *SEL1L-FLAG* constructs (WT, S658P, L709P, P699T or ΔAPH). Quantification of HMW AVP shown below the gel represents the average of three independent experiments, normalized to the loading control HSP90.
Values are shown as mean ± SEM. n.s., not significant; *p < 0.05; **p < 0.01; ***p < 0.001; ****p < 0.0001 using one-way ANOVA with Dunnett's multiple comparisons test.

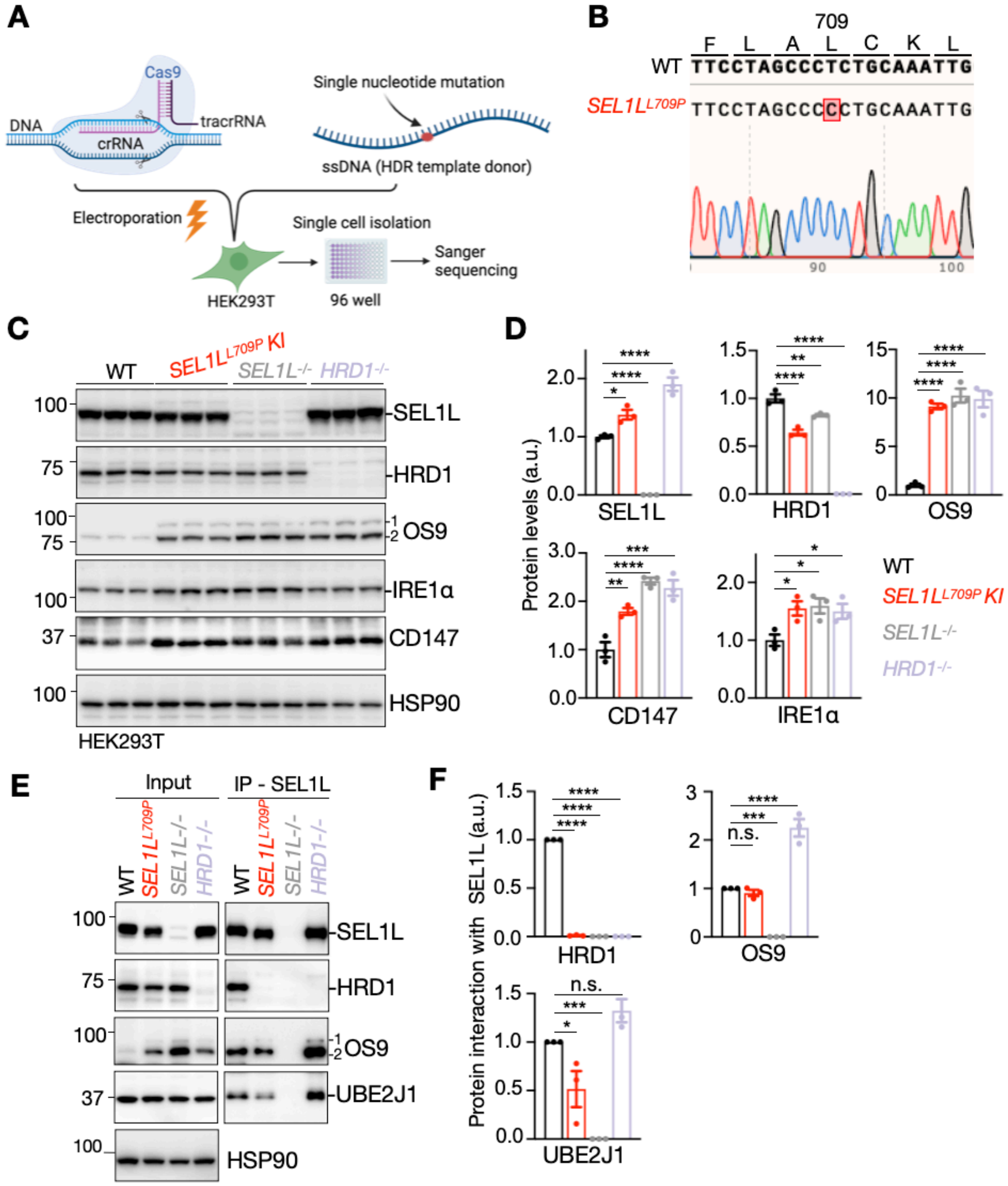


**Figure S5. SEL1L L709P mutation abolishes SEL1L-HRD1 interaction in human cells.**
(A) Schematic of CRISPR/Cas9-mediated KI of the L709P mutation in HEK293T cells.
(B) Sanger sequencing confirmation of biallelic KI cell lines.
(C–D) Immunoblot analysis of ERAD-related components and substrates in WT, *SEL1L*$^{L709P}$, *SEL1L*$^{-/-}$, and *HRD1*$^{-/-}$ HEK293T cells (C). Quantification is shown in (D), normalized to the loading control HSP90; Data represents three independent experiments, with 2 repeats.
(E-F) Co-immunoprecipitation (Co-IP) of endogenous SEL1L in the indicated HEK293T cell lines, followed by immunoblotting for ERAD components (E). The *L709P* mutant shows loss of HRD1 binding but retains interactions with other ERAD factors. Quantification is shown in (F), normalized to SEL1L; HSP90 is the input loading control.
Values, mean ± SEM. n.s., not significant; *p < 0.05; **p < 0.01; ***p < 0.001; ****p < 0.0001 using one-way ANOVA with Dunnett's multiple comparisons test.

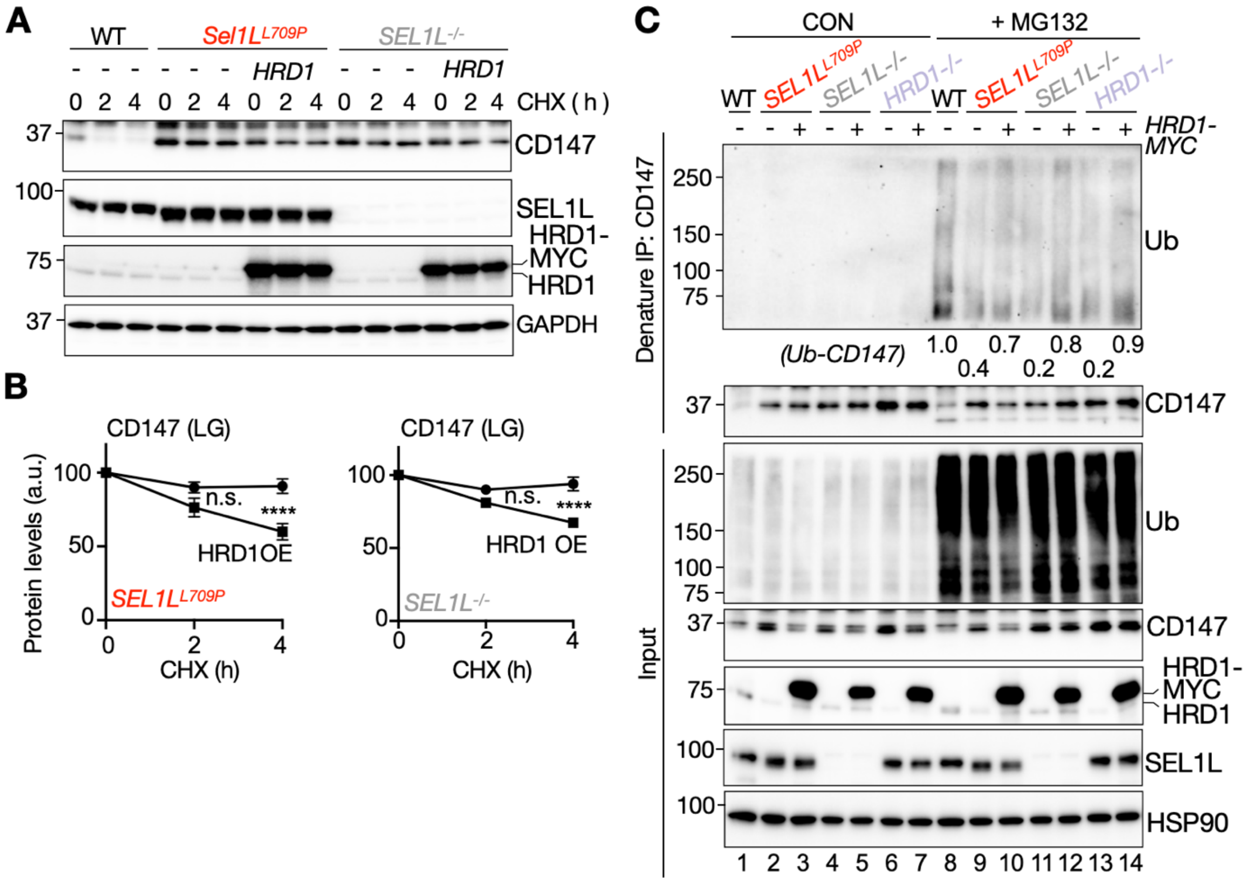


**Figure S6. HRD1 overexpression partially rescues ERAD defects in *SEL1L*$^{L709P}$ cells.**
(A–B) CHX chase analysis of CD147 degradation in WT, SEL1L-/-, and SEL1LL709P cells with or without HRD1-MYC OE (A). Quantification is shown in (B), normalized to GAPDH. Data represents the n = 4 independent experiments, with 2 repeats.
(C) Denaturing IP detecting ubiquitinated CD147 with or without HRD1-MYC OE in WT, *SEL1L*$^{-/-}$, and *SEL1L*$^{L709P}$ cells. Quantification of CD147 ubiquitination shown below the gel and represents the average of three independent experiments.
Values, mean ± SEM. n.s., not significant; ****p < 0.0001 using two-way ANOVA with Tukey's post hoc test (B).
